# USP47 regulates a 53BP1-based complex function in DNA DSB repair

**DOI:** 10.1101/2021.01.27.428282

**Authors:** Doraid T. Sadideen, Baowei Chen, Manal Basili, Robert Hromas, Montaser Shaheen

**Author notes:** Corresponding Author: Montaser Shaheen, University of Arizona, 1515 N. Campbell Avenue, Tucson, AZ 85724.

## Abstract

53BP1 is a key DNA repair protein that regulates double strand break (DSB) repair pathway choice. It assembles a complex, whose loss reverses BRCA1 deficient cells sensitivity to PARP1 inhibitors potentially by antagonizing DSB end resection. 53BP1 is critical for immunoglobulin heavy chain class switch recombination (CSR), that is mediated by the generation and repair of DSBs. Here we identify a deubiquitylase, USP47, as a 53BP1 associated protein that regulates its function in these processes. USP47 loss results in 53BP1 protein instability in a proteasome dependent manner and in reduced 53BP1 ionizing radiation induced foci (IRIF). Like 53BP1, USP47 depletion results in enhanced homologous recombination repair, and defective immunoglobulin CSR. It is also required for resistance to DNA damaging agents. We identify a complex of 53BP1, USP47, Fen1, and Recql1 that regulates a microhomology mediated end joining (MMEJ), a repair pathway that is involved in CSR. Altogether, our findings implicate USP47 in DNA DSB repair at least through regulating a novel 53BP1 associated complex.

## Introduction

DNA double strand breaks (DSBs) are some of the most toxic lesions and if non-or mis-repaired, leads to genome instability and cancer^1^. There are several DSB repair pathways that fall into two major categories: Homologous recombination (HR) and non-homologous end joining (NHEJ)^2^.

TP53 Binding protein 1 (53BP1) is a key DSB repair protein that regulates the repair pathway choice^3^. Thousands of 53BP1 molecules accumulate through specific chromatin interactions, oligomerization and liquid-liquid phase separation, to form Ionization irradiation induced foci (IRIF)^4^ around a single or a cluster of DSBs. 53BP1 recruits a machinery that antagonizes excessive DSB end resection and blocks HR through a pre- and post-resection mechanisms^5^. Large scale Proteomic experiments uncovered many proteins that associate with 53BP1 and likely contribute to the repair process through known and unknown mechanisms^6^. Ubiquitin-associated processes are abundant at the repair sites, and they are required for 53BP1 recruitment, although the functional links of several ubiquitin-pathway proteins keep evolving or remain unknown^7^. 53BP1 was reported to be ubiquitylated at damage sites and an E2 ubiquitin ligase was implicated in its degradation^8^.

USP47 is one of about 100 human deubiquitylases (dubs) that regulate a massive number of ubiquitylation events that involve all cellular functions^8^. USP47 is involved in DNA Base Excision Repair (BER) through interacting with and deubiquitylating the polymerase Pol β^9^. It is phosphorylated at defined residues post IR and etoposide^10^, which suggests involvement in DNA DSB repair. USP47 displays significant homology to USP7, a deubiquitylase that is involved in a plethora of chromatin and DNA damage processes^13^. Through ongoing proteomic and functional screens to identify 53BP1 regulators or effectors, we identified USP47 as a novel 53BP1-asscoiated protein. We show that USP47 is required for 53BP1 stability and recruitment to damage sites and that it is involved in the same biological processes that pertain to 53BP1. Moreover, we identify a complex that harbors both proteins and that is involved in micro-homology mediated end-joining (MMEJ) of DNA DSBs.

## Results

### USP47 associates with 53BP1 and enhances its recruitment/ retention at the damage sites

While significant progress has been made in uncovering 53BP1 biology, new players continue to emerge^3^. In an ongoing siRNA screen of human deubiquitylases’ with 53BP1 IRIF as a readout, we identified BRCA1 associated protein (BAP1) and USP47 as top hits. Details of the screen and BAP1 depletion phenotype will be presented elsewhere. We also did several proteomic pulldowns of 53BP1 complex utilizing matrix-bound antibodies and nanobodies against tagged and native 53BP1 protein followed by mass spectrometry fingerprinting. We identified USP47 in one pulldown in which we used high affinity antibody against the endogenous 53BP1 protein (Figure 1A). USP47 peptides were present only in the 53BP1 pulldown and absent in several control runs that were completed at the same time and in the same conditions. We also pulled down RIF1, a known 53BP1 partner, in the same experiment. We confirmed 53BP1 and USP47 association by IP followed by WB utilizing antibodies against tagged and native proteins (Figure 1B & C). As noted in Figure1, the two proteins association is enhanced after exposing cells to IR suggestive of USP47 involvement in DSBs repair. While USP47 did not form obvious IRIF upon IR exposure (utilizing ab against the endogenous protein), we found that the two protein colocalize on a cluster of DNA DSBs trigged by the nuclease FOK1, (describe more LacI-Fok1 system (Figure 1D).

**Figure 1:**
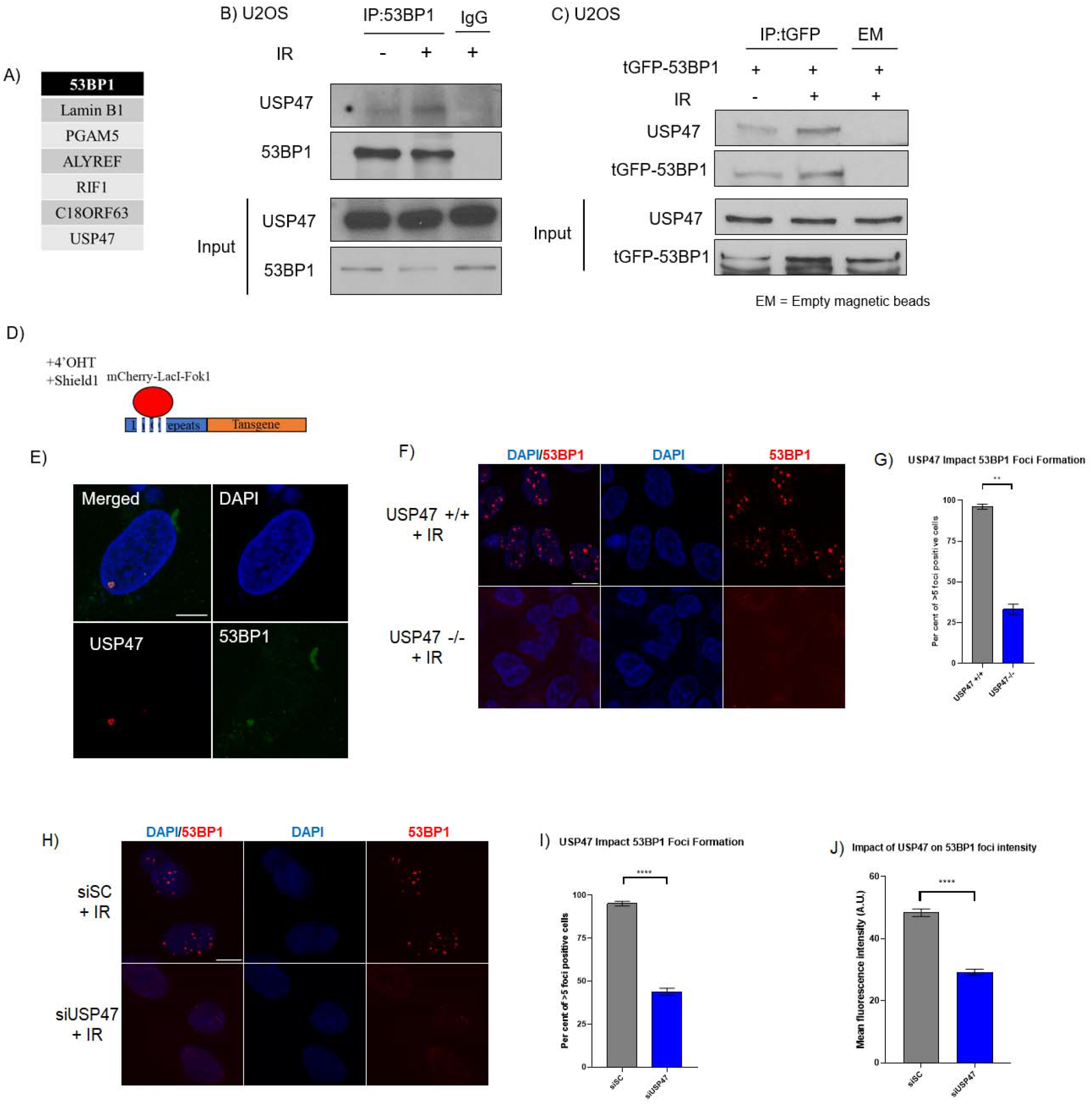
USP47 associates and interacts with 53BP1. **A**. Proteins that appeared in the mass spectrometry fingerprinting of 53BP1 complex immunoprecipitated by a high avidity antibody. None of these peptides were pulled down shows USP47 peptides. **B**. The association between USP47 and 53BP1 increases after ionizing irradiation (IR). **C**. Association between tGFP-53BP1 and USP47 in U2OS. **D**. U2OS reporter cell line transfected with FLAG-USP47 and treated with 4-OHT for 4 hrs to induce DSBs by mCherry-Lac1-FokI. USP47 co-localizes with 53BP1 on DSBs locus. **E**. FOK1 DSB reporter cell line showing c-localization between USP47 and 53BP1. **F**. CRISPR-generated A549 USP47 deficient cells shows decreased 53BP1 IRIF compared to control cells **G**. Histogram quantification of **F. H**. U2OS cells treated with siRNA against USP47 shows similar phenotype of decreased 53BP1 IRIF. **I**. Histogram quantification of **H** showing the decrease in 53BP1 IRIF. **J**. Histogram showing the mean foci intensity. Scale bar is 10μm. P-value ** <0.001, **** <0.0001

As we observed in the screen, 53BP1 recruitment to damage sites was significantly decreased in USP47 depleted cells by CRISPR and siRNA (Figure 1E). Both of 53BP1 IRIF number and focus intensity were impaired in in USP47 depleted cells (Figure 1F & G). The phenotype after siRNA depletion was even more pronounced probably due to the use of different cell lines (U2OS in siRNA vs A549 in CRISPR) and potentially due to adaptation after CRISPR knockdown and establishing a stable line.

### USP47 regulates 53BP1 Stability through deubiquitylation

We hypothesized that USP47 depletion might cause 53BP1 protein instability, and indeed, reducing USP47 protein level by siRNA or CRISPR is associated with a decrease in 53BP1 protein level particularly after IR (Figure 2A, B, C). 53BP1 protein declines faster in USP47-/-compared to USP47+/+ cells after halting new protein synthesis consistent with reduced 53BP1 half-life after IR (Figure 2D). The decrease in 53BP1 protein level in USP47 depleted cells was reversed with the addition of a proteasome inhibitor consistent with a proteasome-mediated degradation (Figure 2E & F). We examined 53BP1 ubiquitylation status in human cell lines after proteasome inhibition by collecting the lysate under denaturing conditions to preserve the ubiquitylated species and we observed enhanced 53BP1 high molecular weight ubiquitylated 53BP1 in USP47 deficient cells (Figure 2G).

**Figure 2:**
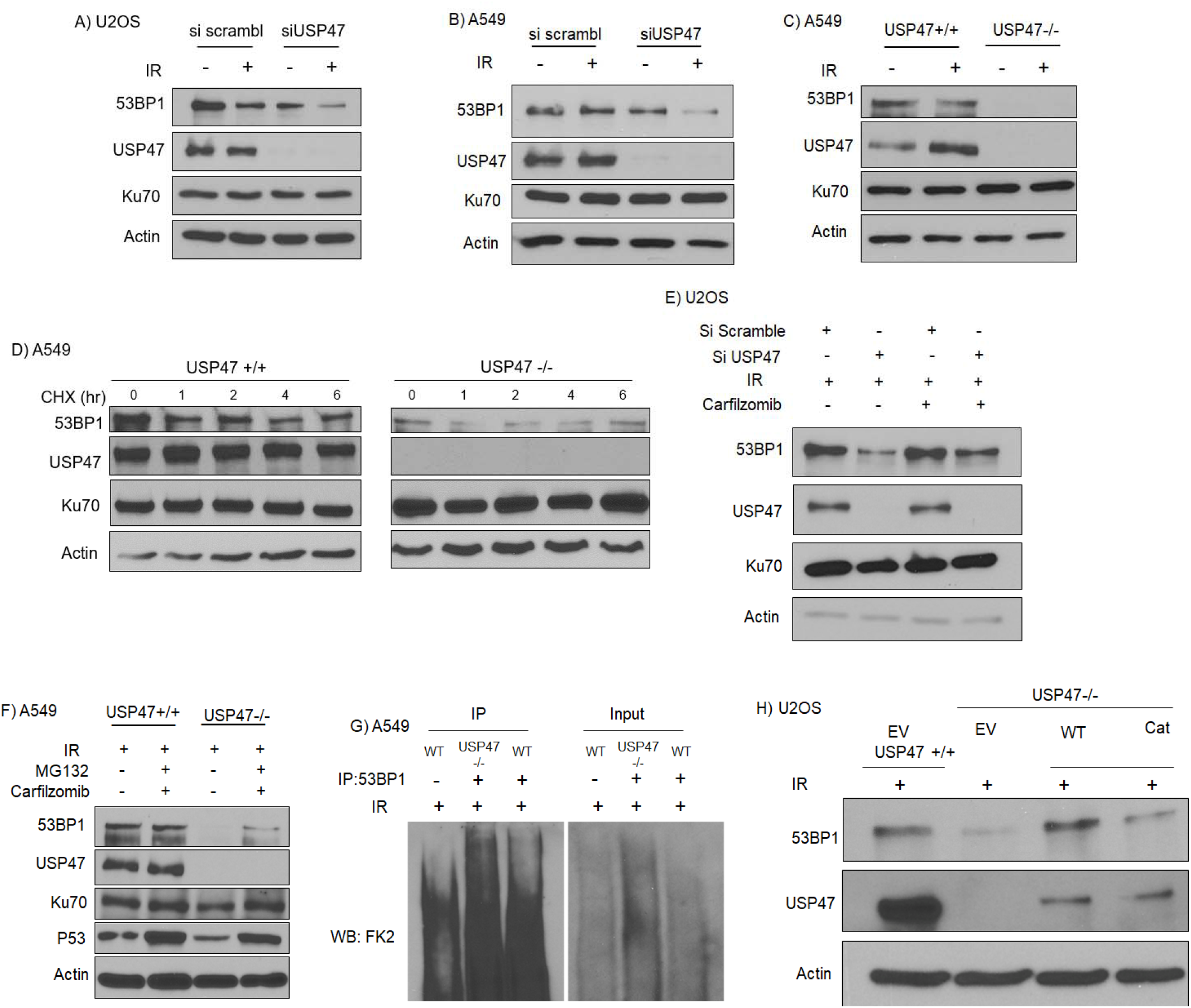
USP47 regulates 53BP1 stability through deubiquitylation. **A-B**. Transient knockdown of USP47 using siRNA in U2OS and A549 cells, respectively, shows 53BP1 instability after IR. **C**. USP47 CRISPR-generated A549 knockout cell line. **D**. A549 proficient and deficient USP47 cells treated with Cycloheximide and collected after IR after indicated time points. **E**. CRISPR-generated USP47 A549 proficient and deficient cell line shows a 53BP1 instability phenotype, which is partially rescued with proteasome inhibitor Carfilzomib. **F**. 53BP1 protein levels are partially rescued in siUSP47 treated cells after proteasome inhibitor treatment. **G**. Enhanced ubiquitylation of 53BP1 in USP47 -/- cells compared to USP47+/+ cells. **H**. expression of wild-type USP47 rescues 53BP1, while catalytically inactive USP47 does not in CRISPR-generated USP47 deficient cells.

To investigate the role of USP47 catalytic activity in the observed phenotypes, we mutated a critical cysteine in position 144 to alanine within the USP protease domain. Wild type USP47 but not catalytically dead mutant was able to reverse 53BP1 protein instability (Figure 2H). Re-expression of wild type USP47 in CRISPR USP47 deficient A549 cells rescued 53BP1 IRIF formation and fluorescence intensity, while catalytically inactive USP47 did not (Figure 3A & B). Altogether, our findings indicate that USP47 protects 53BP1 from proteasome mediated degradation through deubiquitylation that is mediated by its catalytic activity.

**Figure 3:**
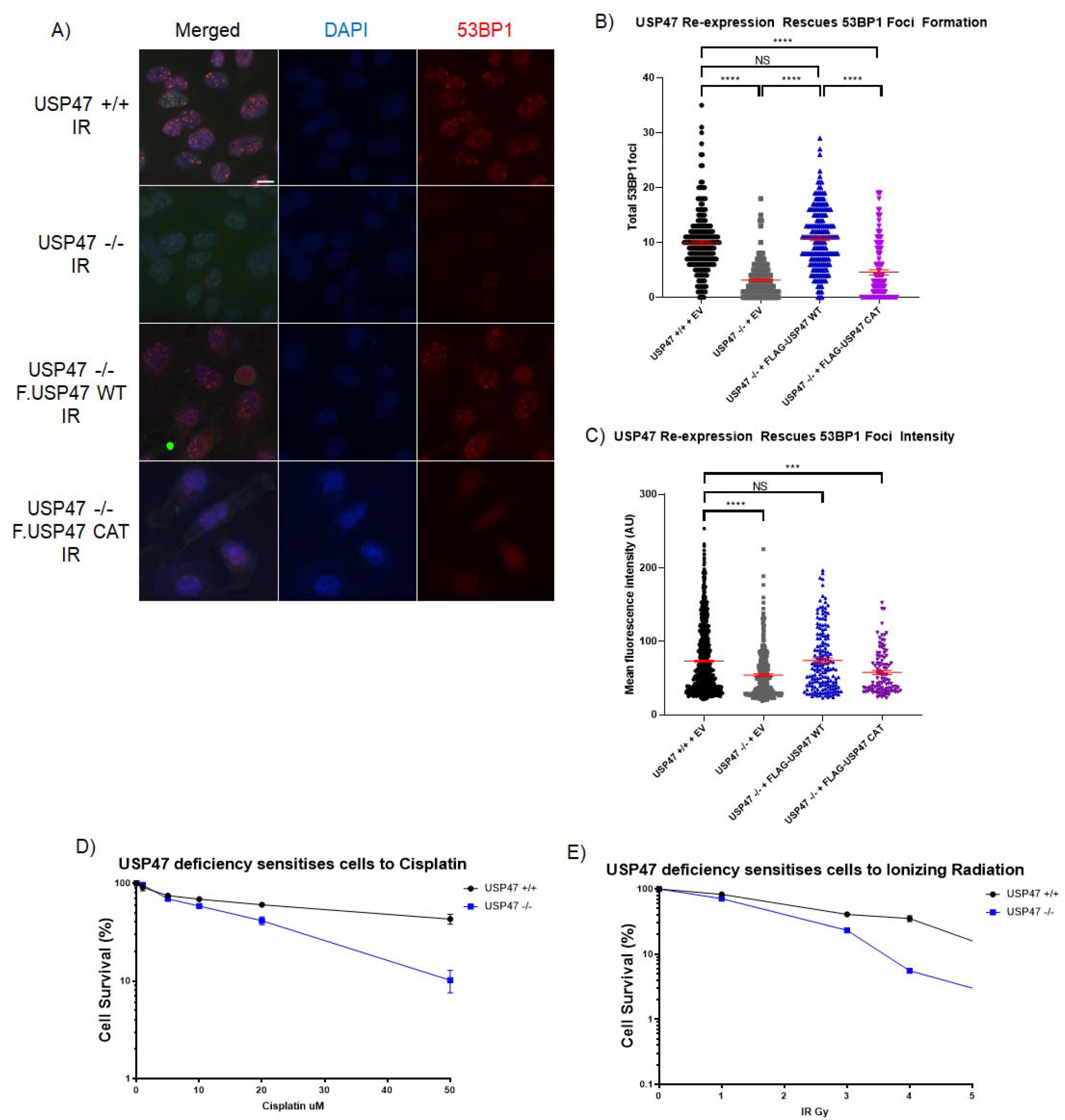
USP47 loss sensitizes cells to IR and cisplatin treatment. **A**. CRISPR-generated A549 USP47 proficient and deficient cells show 53BP1 foci rescue by expression WT but not catalytically inactivate USP47. **B** Quantification of 53BP1 foci formation of photomicrographs in **A. C**. Quantification of 53BP1 mean foci intensity. **D**. USP47 proficient and deficient cells treated with Cisplatin at indicated dosages and left to grow for 14 days. **E**. Similar to **D** except cells exposed to indicated IR doses. Scale bar is 10μm. P-value: <0.0001.

### USP47 depletion mimics 53BP1 loss in DNA damage resistance, impaired immunoglobulin CSR and hyperactive HR repair

53BP1 deficient cells are sensitive to DNA damaging agents that generate DSBs^4^. Similarly, we observed higher sensitivity of cells depleted of USP47 to IR and cisplatin compared to control isogenic cells (Figure 3C & D).

A primary function of 53BP1 is to facilitate immunoglobulin heavy chain class switch recombination (CSR)^14,15^, a process that involves generating DSBs at the Immunoglobulin switch regions and that relies on classic and alternative NHEJ and 53BP1 deficiency leads to a dramatic CSR defect. Given USP47 requirement to maintain 53BP1 stability and presence at DSBs, we investigated its contribution to CSR. To this end, we utilized murine lymphocytic cells (CH12F3) that switch immunoglobulins upon cytokine stimulation^16^. We depleted USP47 by lentivirus shRNA, and we identified significant impairment in CSR although not to the same profound impairment that is seen with 53BP1 total loss by CRISPR (Figure 4A& B). 53bp1 blocks excessive resection at DNA DSB ends, and its depletion results in hyper-resection and enhanced DSB repair by HR at pre- and post-resection levels^3,5^. To investigate USP47 effect on HR, we utilized a widely used system (DR-GFP) that measures HR repair of DSB induced by the rare cutting restriction enzyme I-sce1. We found that USP47 depletion enhanced HR like the phenotype seen after 53BP1 depletion (Figure 4C & D).

**Figure 4:**
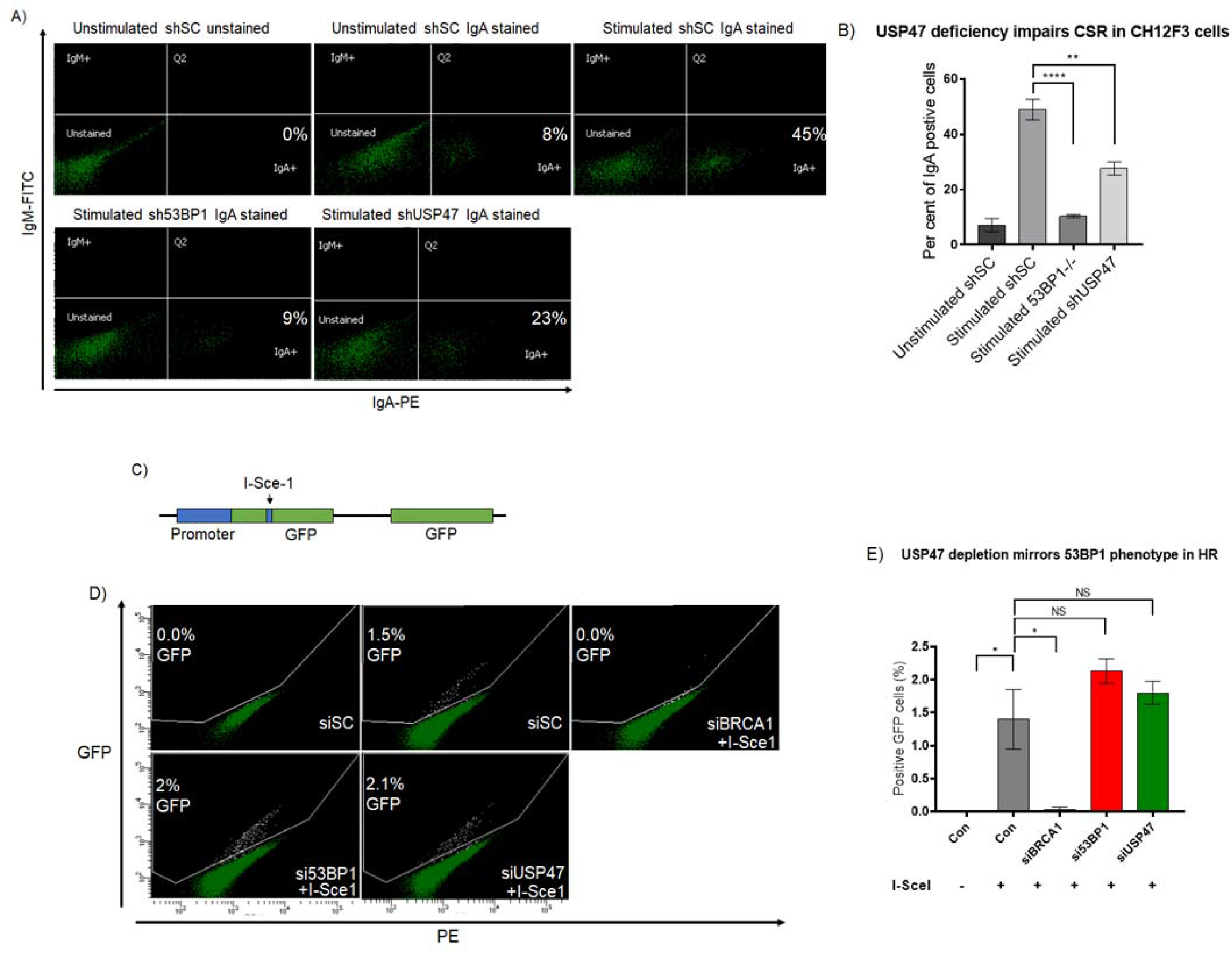
USP47 loss impairs immunoglobulin CSR and hyperactive HR repair. **A**. Flowcytometry showing percentage of cells that switched from IgM to IgA after knockdown of indicated genes. **B**. Histogram quantification of A. **C**. HR substrate schematic. **D**. flowcytometry gating on GFP-positive cells after knockdown of indicated genes. **E**. Quantification of **D**.

### USP47 is part of a base excision repair (BER) complex that associates with 53BP1 and promotes micro-homology mediated joining (MMEJ)

USP47 interacts with and regulates DNA polymerase B (Pol B) function in BER^9^. Pol B and other players in the BER (such as PARP1, DNA Ligase III and FEN) have been reported to be involved in DSB repair through MMEJ^17,18,19^. The DNA helicase RECQL1 interacts and enhances FEN1 nuclease function^20^ and RECQL1 is involved in long patch BER as well^21^. We noticed that both of Fen1 and RECQL1 are associated with USP47^22^ and 53BP1^5,6^ in large scale proteomic analysis. FEN1 is involved in long patch BER^23^ and in MMEJ^18,19^. While 53BP1 suppresses HR it has not been directly implicated in class NHEJ, although it was reported to be involved in MMEJ^24^. Based on the above functional and physical connections and clues, we envisioned the presence of a complex (53BP1-USP47-FEN1-RECQL1) that facilitates MMEJ of DSBs. To prove that we first performed reciprocal Immunoprecipitation experiments and showed that these proteins associate with each other (Figure5). The association between USP47 with RECQL1 increased after IR (Figure5A). We examined this complex contribution to MMEJ by depleting each individual protein in a cell line that harbors a MMEJ substrate integrated into its genome (U2OS EJ-2). We found that depleting each of these four proteins results in defective MMEJ (Figure 5E & F). FEN1 and RECQL1 involvement in MMEJ fits well with their nuclease and helicase activities that are likely needed to process DNA flaps generated at the DNA termini and to facilitate ligation^19,25^. In the same context DNA ligase III, a well-established MMEJ protein^18^, comes down with heavy peptide load within the 53BP1 complex (our own unshown mass spec data, also observed in Refs 5,6). We confirmed that Lig III associates with 53BP1 by IP (Figure 5D), which strengthens the evidence of 53BP1 involvement in MMEJ.

**Figure 5:**
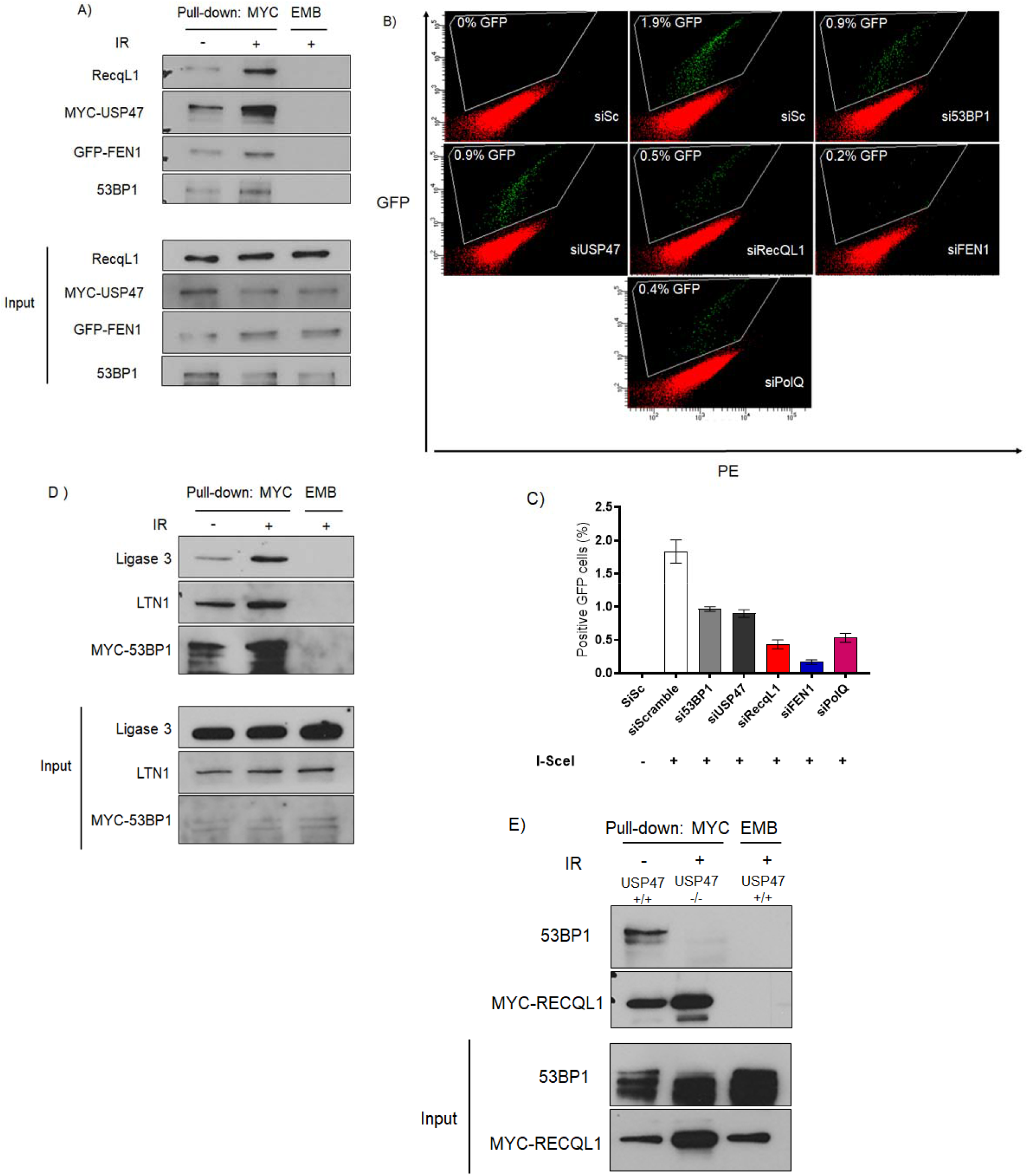
USP47-53BP1 complex that regulates MMEJ. **A**. Myc-USP47 pulls down RECQL1, FEN1, and 53BP1. **B**. GFP-positive cells in MMEJ reporter show impaired assay when each of this complex proteins are depleted. **C**. Histogram of **B. D**. MYC-53BP1 pull-down shows interaction with LTN1 and ligase 3. **E**. Myc-RECQL1 pulls down 53BP1 and the association is decreased in USP47 deficient cells.

We asked if USP47 through its association with FEN1 and RECQL1 might have other activities in this complex besides stabilizing 53BP1. We investigated if USP47 depletion impacts FEN1 or RECQL1 ubiquitylation or interaction with 53BP1. There was no significant effect on their protein levels (Supplement Figure), but RECQL1 association with 53BP1 was lost in USP47 deficient cells (Figure 5B), and this lost association was more than expected due to reduced 53BP1 protein level. We hypothesized that USP47 might regulate RECQL1 through deubiquitylation. When we investigated RECQL1 ubiquitylation, we identified two ubiquitin bands representing mono- and di-ubiquitin RECQL1, and we observed enhanced RECQL1 ubiquitylation in USP47-/- cells vs control (Figure 6A), which potentially can impede its interaction with 53BP1.

**Figure 6:**
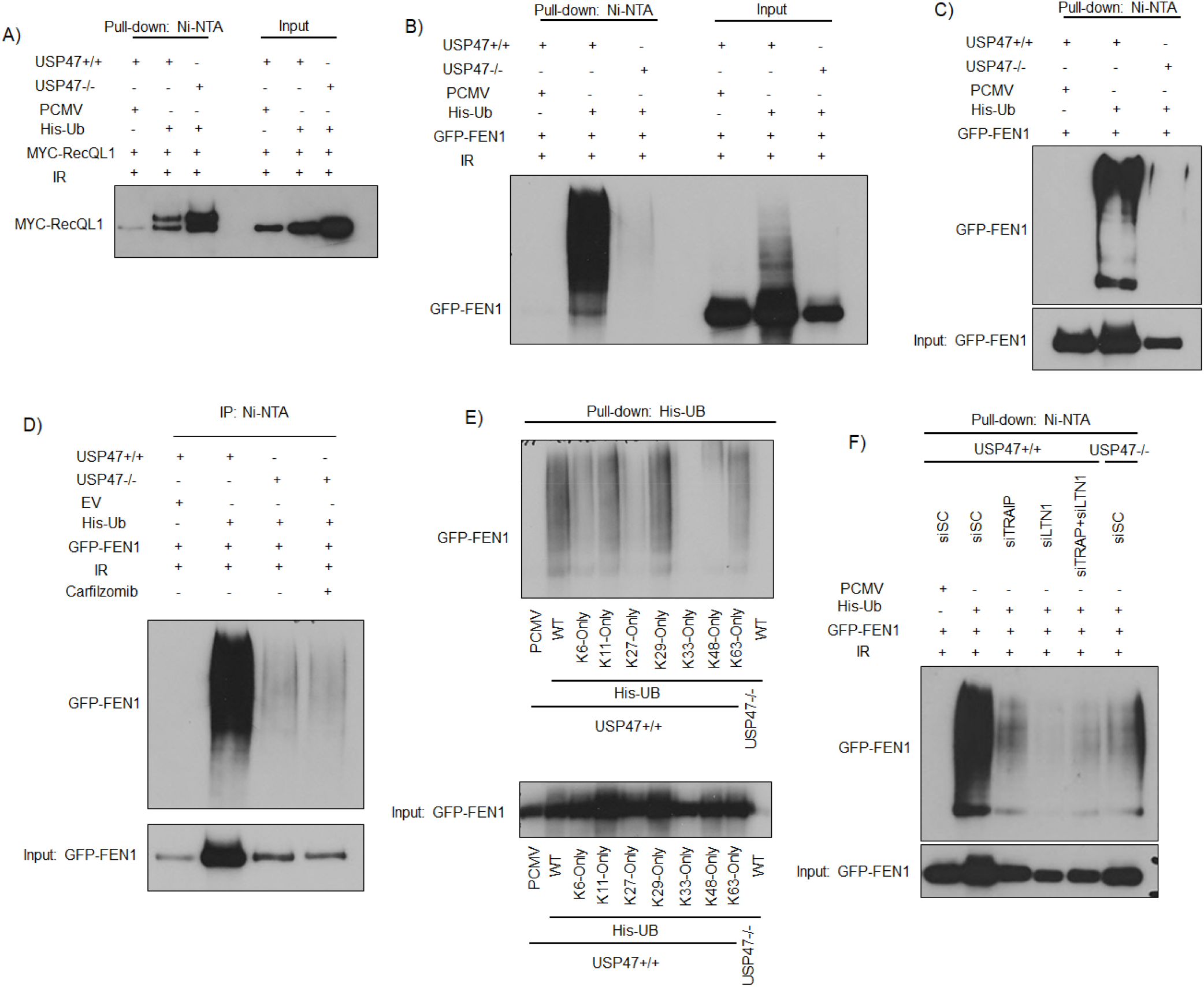
RECQL1 & FEN1 ubiquitylation in USP47 depleted cells. **A**. RECQL1 ubiquitylation is enhanced in USP47-/- cells. **B**. FEN1 is heavily ubiquitylated in USP47+/+ cells and this is markedly decreased in USP47-/- cells. **C**. FEN1 is ubiquitylated in a SteadyState condition without inducing DNA damage. **D**. Proteasome inhibitor treatment does not change the hypo-ubiquitylated state of FEN1 in USP47 deficient cells. **E**. His-UB WT and Lysine mutants where only the indicated lysine is present, shows FEN1 is ubiquitylated on K63 and K48. **F**. Each of TRAIP and LTN1 E3 ligases are needed for full FEN1 ubiquitylation.

On the contrary, we found that FEN1 is heavily ubiquitylated after IR in USP47+/+ A549 cells and this ubiquitylation is markedly decreased in USP47-/- cells (Figure 6B). FEN1 ubiquitylation was also observed at baseline and without subjection the cells to DNA damage (Figure 6C). This paradoxical phenotype of reduced ubiquitylation after a deubiquitylase (USP47) depletion suggests that FEN1 may be subjected to different layers of ubiquitylation that may regulate its function or that USP47 may indirectly impact FEN1 ubiquitylation through other factors. Inhibiting the proteasome did not reverse the decrease in FEN1 ubiquitylation in USP47-/- cells ruling out proteasome mediated protein instability as an explanation for the observed phenotype (Figure 6D). We mutated the ubiquitin plasmid in all the lysine residues that mediate ubiquitin chain formation and identified that FEN1 is ubiquitylated utilizing chains that binds through K6, K29 and K63 (Figure 6 E). We aimed to identify the ubiquitin E3 ligase that mediates FEN1 ubiquitylation. By searching publicly available proteomic data, we found that FEN1 is associated with a couple of ubiquitin ligases, namely TRAIP and Listerin (LTN1)^12,22^. Depleting each of these two E3 ligases resulted in marked reduction in FEN1 ubiquitylation (Figure 6F). TRAIP has been implicated in DNA DSB repair^26^, and LTN1 peptides can be identified within 53BP1 interactome complex^6^. We confirmed that LTN1 is pulled down by 53BP1 (Figure 5D). We conclude that FEN1 is ubiquitylated by TRAIP and LTN1and that USP47 is required for its ubiquitylation.

## Discussion

53BP1 has emerged as a key regulator of DSB repair by assembling a machinery that protects DSB ends from excessive resection and antagonizes BRCA1 function at DSB level. Here we show that USP47 is critical for 53BP1 stability, focus formation, and activity in DSB repair, through its catalytic activity. Other lines evidence implicate USP47 in DNA DSB repair and lend support to our model as follows. USP47 is phosphorylated at defined residues post IR and etoposide^10^. MDC1, a critical DSB repair protein and a 53BP1 partner^11^, is also associated with USP47 based on a large proteomic study^12^. USP47 was previously implicated in another DNA repair pathway (BER) by stabilizing the polymerase Pol β^9^. A few BER proteins are recruited to DSB to participate in DSB repair by MMEJ^25^. We show here that USP47 is also involved MMEJ. This DSB repair pathway is important for immunoglobulin CSR^27^, a process that heavily relies on 53BP1^14,15^. Thus, a role for 53BP1 in MMEJ is not surprising from a functional perspective. Another report implicated 53BP1 in MMEJ using a different MMEJ assay and it revealed that 53BP1 promotes deletions of moderate length at repair junctions that could hint to the involvement of a nucleases such as FEN1^24^. 53BP1 inhibits extensive resection mediated by other nucleases such as DNA2 and EXO1 potentially through the Shieldin complex likely to suppress other mutagenic repair pathways such as single strand annealing (SSA)^28^, and while 53BP1 depletion enhances HR, it has not been reported to be directly involved in classical NHEJ and thus a physical involvement as a scaffold in at least one repair pathway such as MMEJ is plausible.

We implicated RECQL1, which is involved in BER^21^ and in the repair of DNA DSBs^29^, in MMEJ. RECQL1 is also pulled down within the Shieldin complex^30^. We show that RECQL1 undergoes ubiquitylation that likely regulates its DNA damage function. Future efforts will likely identify the ubiquitin ligases that target RECQL1 and the role of this ubiquitylation in its function. We also show that FEN1 is ubiquitylated in the steady state condition and we describe two ubiquitin ligases that regulates FEN1 ubiquitylation. TRAIP1 has been implicated in ICL repair and in variable ubiquitylation of the GMC helicase to regulate its unloading at the damage site. LTN1 was identified in the 53BP1 complex^6^, and here we implicate it in FEN1 ubiquitylation. We show that mixed ubiquitin chains assemble on FEN1 and likely regulates either its localization, function or interactions with other proteins. The mechanism of FEN1 ubiquitin smear attenuation in USP47-/-is likely indirect will be investigated in future efforts.

Polymerase theta (PolQ), is a primary mediator of a sub-pathway of MMEJ (Theta mediated End Joining or TMEJ)^31^. Notably, 53BP1 and its pathway partners display significant synthetic lethality with PolQ^32^ suggesting the two protein regulates complementary and not identical pathways of repair. Recent data revealed that PolQ loss produces a repair product phenotype that is shared with the loss of another complex (Rad17 and RFC proteins) rather than the established MMEJ repair factors such as LigIII or FEN1^33^.

53BP1 regulation by ubiquitin and ubiquitin-like proteins is likely extensive given the number of ubiquitin and sumo-associated proteins that are discovered in proximity to 53BP1^6^. Future research will likely identify which ubiquitin ligase(s) destines 53BP1 to the proteasome and will further shed light into other hidden substrates/ functions of USP47.

## Materials and Methods

### Cell Lines, CRISPR and Gene Silencing

Human HEK293 (ATCC, CRL-1573), A549 (ATCC, CRM-CCL-185) and U2OS (ATCC, HTB-96) cells were maintained in Dulbecco’s Modified Eagle Medium (DMEM, Corning) supplemented with 5% heat-inactivated fetal bovine serum (FBS) (Genesee Scientific) in a 5% CO2 incubator at 37°C. CH12F3-2 and CH12F3-2 53BP1-/-murine B cell lymphoma cells^17,18^ were cultured in RPMI 1640 medium supplemented with 15% FBS, 50nM 2-Mercaptoethanol (BME), 5% NCTC-109 (Life Technologies)and 20 mM HEPES (Corning).

Short-term depletion of USP47 expression was achieved by treating cells with siRNA USP47 (5’-GAA AGC AAA UGA AGG GAA AUU-3’ or 5’-AGA UAA AGA UGG UGA ACA AUU-3’) from Dharmacon and Lipofectamine RNAiMAX transfecting reagent twice according to the manufacturer’s instruction (Invitrogen). All siRNAs were purchased from Dharmacon unless otherwise stated. Other siRNA used in the study are:

53BP1: Sense: 5’-GAA CAG AAG UAG AAA GAA AUU-3’

LTN1: Sense: 5’-CGA UAU CCU UGG UGA GAA AUU-3’

TRAIP: Sense: 5’-CAG CAU GGU UAC UAC GAA AUU-3’

Scramble from Qiagen Cat No.: 1027310: Sense: 5’-UUC UCC GAA CGU GUC ACG UdT dT-3’

To generate CRISPR deficient USP47 cell line transfection was performed according to IDT’s instruction manual with modification. Briefly, using reverse transfection, A549 cells were trypsinized and plated 5×10^5 cells per well in 6-well plate mixed with 800µL of RNP complex reverse transfection and crRNA sequence are 5’-ACTGTCATCACGGAGTACAA-3’ and 5’-GCAATATAGTAGAAACGAGT-3’ or proprietary non-targeting sequence for control. After 24hrs, the complex was removed and fresh DMEM supplemented with 10% FBS was added. The cells were trypsinized 48hrs later and sorted through FACS in 96-well plates for colonies. Two week later, the 96-well plates were screened for homozygous knockout of USP47.

### Mass Spectrometry

U2OS cells expressing different GFP-tagged FEN1, tGFP-tagged or MYC-tagged USP47 or 53BP1 proteins with a proper control were collected from 10x 100-mm plates and lysed in IP buffer (50mM HEPES-KOH, pH8.0, 100mM KCl, 2mM EDTA, 0.1% NP-40, 10% glycerol, complimented with mixture of protease and phosphatase inhibitors). Cells were sonicated using QSonica Q125 at 60% Hz for 10 seconds (QSonica, CT, USA), and centrifuged at max speed for 1 hour. Lysates were affinity-purified using GFP-Trap, tGFP-Trap or MYC-Trap Nano magnetic beads (ChromoTek). Mass spectrometry analysis was carried at Taplin Mass Spectrometry Facility (Harvard Medical School, MA, USA). All immunoprecipitations were performed in three biological replicates.

### Reagents and Plasmids

The proteasome inhibitors MG132 and Carfilzomib were purchased from Sigma-Aldrich and TLC Pharmaceutical Standards Ltd., respectively. Protein synthesis inhibitor cycloheximide was purchased from Sigma-Aldrich. USP47 catalytic domain was mutated by changing cysteine 177 into alanine (C177A) using the sense primer (5’-gtttgcaaaaggctattcaaataggcagtcattgcttggtttactagtcc-3’) using Quickchange Lightning Site-directed Mutagenesis Kit from Agilent technologies. MYC-RECQL1 was purchased from Origene (RC200427), GFP-FEN1 was a gift from Professor Muramatsu (National Institute of Infectious Disease, Japan) 1A citation. His-Ub WT and mutant plasmids were a gift from Professor Rape (UC Berkeley, USA) 1B. pRK5-HA-Ubiquitin-WT and mutants were a gift from Ted Dawson (Johns Hopkins, USA) 1C.

### DNA Repair Assays

For DR- and EJ-2-reporter assays, U2OS cells harboring the respective GFP expression cassette were transfected with siRNA (indicated in figure) for two days. After 48 hours, cells were infected with lentiviral BFP-I-SceI or lenti empty vector (BFP) or transfected with I-SceI plasmid. After 48 h, cells were collected, washed and suspended in PBS. GFP-positive cells were analyzed by FACS.

### Western Blotting, Immunoprecipitation and Antibodies

Cell were washed with PBS (137mM NaCl, 2.7 mM KCl, 10mM Na_2_HPO_4_, 1.8mM KH_2_PO_4_) then collected 4hrs after irradiation at a single dose of 6Gy using a Cobalt-60 Source Gamma Irradiator 72hrs after siRNA exposure in lysis buffer (150mM NaCl, 50mM Tris-HCl pH 8, 1mM Dithiothreitol (DTT), 0.5% NP-40, with protease and phosphatase inhibitors (Sigma)), sonicated for 10 seconds twice, and centrifuged at 4°C at 14Kx g for 1h. Protein quantification performed using Bradford Protein Assay (Bio-Rad), and 20µg samples were resolved on a 7% SDS-PAGE gels, transferred to nitrocellulose or PVDF membranes, blocked with 5% non-fat milk or 2% BSA then incubated with primary antibodies overnight. Horseradish peroxidase secondary antibodies and enhanced chemiluminescence (ECL) substrate from Santa Cruz Biotechnology were used to detect antigen-antigen interaction.

Cells were collected with IP lysis buffer (150mM NaCl, 50mM Tris-HCl pH 8, 1mM EDTA, 1mM DTT, 0.5% NP-40, with protease and phosphatase inhibitors (Sigma)), sonicated for 10 seconds twice at 60Hz, and centrifuged at 4°C 14Kx g for 1hr. Samples were pre-cleared using protein G magnetic beads (Dynabeads, Invitrogen). The lysate then incubated overnight with beads conjugated with corresponding antibodies. The beads were washed four times with IP lysis buffer and four times with PBS then boiled in Laemmli sample buffer.

Antibodies include rabbit anti-53BP1 (ab36823, Abcam and NB100-304, Novus Biologicals), rabbit anti-USP47 (A301-048A, Bethyl Laboratories), mouse anti-β-actin (A2228, Sigma), mouse anti-FK2 (04-263, Sigma), mouse anti-TurboGFP (TA150041, Origene), mouse anti-FLAG M2 (F1804, Sigma), mouse anti-GFP (sc-9996, Santa Cruz), mouse anti-FEN1 (2746S, Cell Signaling), mouse anti-HA (2367S, Cell Signaling), mouse anti-RECQL1 (sc-166388, Santa Cruz) and mouse anti-MYC (2276S, Cell Signaling).

### Immunofluorescence, Confocal Microscopy and Analysis

Cells grown on coverslips were exposed to 1Gy IR. After 4hrs, the cells were fixed for 10 minutes in 4% formaldehyde and sucrose at room temperature. Then permeabilized for 10 minutes in 0.5% Triton-X in PBS and blocked for 1hr in 4% BSA in PBS at room temperature. The cells were washed 3x with PBS before every step. The indicated primaries were diluted in 1% BSA in PBS, while Alexa Fluor 488 and 568 secondary antibodies were diluted in PBS. Hoechst 33342 (1μg/mL) was added on the cells with secondary and incubated at room temperature for 1hr in a dark room. Cells were washed 3x with PBS and mounted using ProLong™ Gold Antifade (Thermofisher, P36934).

Images were visualized on a Leica SP5-II confocal microscope using oil-immersion 63x objective at the University of Arizona Cancer Center. Sequential imaging using a multi-channel utilizing the following laser settings and detectors: 405nm Diode (blue), 488nm Argon (25% laser power, green) and HeNe 543nm (red). Scale bar is for all images is 10μm unless stated otherwise.

Images were exported as non-compressed TIFFs using LAS X software and analyzed using NIS Elements Advanced Research Software (Nikon, NY, USA). An image analysis pipeline was applied to every picture, which segments the cell nuclei by region of interest (ROI). A threshold for speckle (focus) size was applied to each γ-H2AX (0.2μm^2^) and 53BP1 (0.68μm^2^) and foci were counted and compiled by the software.

### Clonogenic Assay

CRISPR-generated USP47 cells were plated at approximately 500 cells per well in 6-well plate or 60mm dishes. The following day, cells were incubated with cisplatin for 2hrs at indicated concentrations then washed three times with PBS and fresh medium was add to the wells or exposed to IR at indicated dosages and left to grow. After 14 days, colonies (defined as >50 cells) were fixed and stained with 1% Crystal violet solution (Sigma).

### In Vivo Deubiquitylation

For *in vivo* deubiquitylation of 53BP1 by USP47, A549 CRISPR-generated USP47 proficient and deficient cells were used to overexpress tGFP-53BP1 using Lipofectamine 2000 according to manufacturer’s instructions. 48hrs after transfection, cells were exposed to 6Gy IR dose and treated with 500uM Carfilzomib for 4hrs. Cells were collected in lysis buffer (150mM NaCl, 50mM Tris-HCl pH 8, 1mM DTT, 0.5% NP-40, with protease and phosphatase inhibitors (Sigma).After 1 h, the reaction was stopped by boiling in SDS-based loading buffer, and the ubiquitylated species was detected by anti-ubiquitin chain FK2 antibody.

### Class switch recombination assay

CH12F3-2 and CH12F3-2 53BP1-/-murine B cell lymphoma cells were treated with either shScramble or shUSP47. For induction of CSR, CH12F3-2 cells were incubated in culture medium supplemented with10ng/mL IL-4 (R&D Systems #404-ML-050), 1ng/mL TGFβ (R&D Systems #7666-MB-005) and 1μg/mLanti-CD40 antibody (Thermo Fisher#16-0401-86) for 48hrs. Anti-IgA-PE was used to stain the cells and subsequently PE signal analyzed on BD FACS Canto II machine.

### Statistical analysis

All results are from, at least, three independent experiments. Student’s t-test was used to compare two groups, while one-way analysis of variance (ANOVA) for comparing more than two groups. Graphs were generated using GraphPad Prism 8.0.2 on Microsoft Windows 10 (La Jolla, CA, USA). P-value less than 0.05 will be considered statistically significant.

## References

1. Kass EM, Moynahan ME, Jasin M. When Genome Maintenance Goes Badly Awry. Mol Cell. 2016 Jun 2;62(5):777–87.

2. Hustedt N, Durocher D. The control of DNA repair by the cell cycle. Nat Cell Biol. 2016 Dec 23;19(1):1–9.

3. Mirman Z, de Lange T. 53BP1: a DSB escort. Genes Dev. 2020 Jan 1;34(1-2):7–23

4. Wilson MD, Durocher D. Reading chromatin signatures after DNA double-strand break Philos Trans R Soc Lond B Biol Sci. 2017 Oct 5;372(1731):20160280

5. Callen E, Zong D, Wu W, et al. 53BP1 Enforces Distinct Pre- and Post-resection Blocks on Homologous Recombination. Mol Cell. 2020 Jan 2;77(1):26-38.e7.

6. Gupta R, Somyajit K, Narita T, et al. DNA Repair Network Analysis Reveals Shieldin as a Key Regulator of NHEJ and PARP Inhibitor Sensitivity. Cell. 2018 May 3;173(4):972-988.e23

7. Jackson SP, Durocher D. Regulation of DNA damage responses by ubiquitin and SUMO. Mol Cell. 2013 Mar 7;49(5):795–807

8. Akimov V, Barrio-Hernandez I, Hansen SVF, et al. UbiSite approach for comprehensive mapping of lysine and N-terminal ubiquitination sites Nat Struct Mol Biol. 2018 Jul;25(7):631–640

9. Parsons JL, Dianova II, Khoronenkova SV, et al. USP47 is a deubiquitylating enzyme that regulates base excision repair by controlling steady-state levels of DNA polymerase β. Mol Cell. 2011 Mar 4;41(5):609–15.

10. Beli P, Lukashchuk N, Wagner SA, et al: Proteomic investigations reveal a role for RNA processing factor THRAP3 in the DNA damage response. Mol Cell. 2012 Apr 27;46(2):212–25

11. Eliezer Y, Argaman L, Rhie A, Doherty AJ, Goldberg M. The direct interaction between 53BP1 and MDC1 is required for the recruitment of 53BP1 to sites of damage. J Biol Chem. 2009 Jan 2;284(1):426–435

12. Chweppe DK, Huttlin EL, Harper JW, Gygi SP. BioPlex Display: An Interactive Suite for Large-Scale AP-MS Protein-Protein Interaction Data. J Proteome Res. 2018 Jan 5;17(1):722–726

13. Valles GJ, Bezsonova I, Woodgate R, et al. USP7 Is a Master Regulator of Genome Stability. Front Cell Dev Biol. 2020 Aug 5;8:717

14. Manis JP, Morales JC, Xia Z, et al. 53BP1 links DNA damage-response pathways to immunoglobulin heavy chain class-switch recombination. Nat Immunol. 2004 May;5(5):481–7

15. Ward IM, Reina-San-Martin B, Olaru A, Minn K, et al. 53BP1 is required for class switch recombination. J Cell Biol. 2004 May 24;165(4):459–64.

16. Nakamura M., Kondo K., Sugai M., Nazarea M., Imamura S., Honjo T. (1996) High frequency class switching of an IgM+ B lymphoma clone CH12F3 to IgA+ cellsInt. Immunol., 8 pp. 193–201.

17. Ray S, Breuer G, DeVeaux M, Zelterman D, Bindra R, Sweasy JB. DNA polymerase beta participates in DNA End-joining. Nucleic Acids Res. 2018 Jan 9;46(1):242–255

18. Soni A, Siemann M, Grabos M, Murmann T, Pantelias GE, Iliakis G. Involvement of poly(ADP-ribose) polymerase-1 and XRCC1/DNA ligase III in an alternative route for DNA double-strand breaks rejoining. Nucleic Acids Res. 2014 Jun;42(10):6380–92.

19. Mengwasser KE, Adeyemi RO, Leng Y, et al. Genetic Screens Reveal FEN1 and APEX2 as BRCA2 Synthetic Lethal Targets. Mol Cell. 2019 Mar 7;73(5):885-899.e6

20. Sami F, Lu X, Parvathaneni S, Roy R, Gary RK, Sharma S. RECQ1 interacts with FEN-1 and promotes binding of FEN-1 to telomeric chromatin. Biochem J. 2015 Jun 1;468(2):227–44.

21. Woodrick J, Gupta S, Camacho S, et al. A new sub-pathway of long-patch base excision repair involving 5′ gap formation. EMBO J. 2017 Jun 1;36(11):1605–1622.

22. Huttlin EL, Bruckner RJ, Paulo JA, et al. Architecture of the human interactome defines protein communities and disease networks. Nature. 2017 May 25;545(7655):505–509.

23. Klungland A, Lindahl T. Second pathway for completion of human DNA base excision-repair: reconstitution with purified proteins and requirement for DNase IV (FEN1). EMBO J. 1997 Jun 2;16(11):3341–8.

24. Xiong X, Du Z, Wang Y, Feng Z, Fan P, Yan C, Willers H, Zhang J. 53BP1 promotes microhomology-mediated end-joining in G1-phase cells. Nucleic Acids Res. 2015 Feb 18;43(3):1659–70.

25. Frit P, Barboule N, Yuan Y, Gomez D, Calsou P. Alternative end-joining pathway(s): bricolage at DNA breaks. DNA Repair (Amst). 2014 May;17:81–97.

26. Wu RA, Semlow DR, Kamimae-Lanning AN,et al. TRAIP is a master regulator of DNA interstrand crosslink repair. Nature. 2019 Mar;567(7747):267–272.

27. Yan CT, Boboila C, Souza EK, et al. IgH class switching and translocations use a robust non-classical end-joining pathway. Nature. 2007 Sep 27;449(7161):478–82

28. Ochs F, Somyajit K, Altmeyer M, et al. 53BP1 fosters fidelity of homology-directed DNA repair. Nat Struct Mol Biol. 2016 Aug;23(8):714–21

29. Lu H, Davis AJ. Human RecQ Helicases in DNA Double-Strand Break Repair. Front Cell Dev Biol. 2021 Feb; 9:640755.

30. Findlay S, Heath J, Luo VM, et al. SHLD2/FAM35A co-operates with REV7 to coordinate DNA double-strand break repair pathway choice. EMBO J. 2018 Sep 14;37(18):e100158

31. Ramsden DA, Carvajal-Garcia J, Gupta GP. Mechanism, cellular functions and cancer roles of polymerase-theta-mediated DNA end joining. Nat Rev Mol Cell Biol. 2021 Sep 14

32. Feng W, Simpson DA, Carvajal-Garcia J, Price BA, Kumar RJ, Mose LE, Wood RD, Rashid N, Purvis JE, Parker JS, Ramsden DA, Gupta GP. Genetic determinants of cellular addiction to DNA polymerase theta. Nat Commun. 2019 Sep 19;10(1):4286

33. Hussmann JA, Ling J, Ravisankar P, Yan J, et L. Mapping the genetic landscape of DNA double-strand break repair. Cell. 2021 Oct 28;184(22):5653–5669

